# Single-cell spatial atlas of tertiary lymphoid structures in ovarian cancer

**DOI:** 10.1101/2023.05.16.540946

**Authors:** Joona Sarkkinen, Ada Junquera, Ella Anttila, Angela Szabo, Fernando Perez, Inga-Maria Launonen, Anna Laury, Julia Casado, Eliisa Kekäläinen, Anniina Färkkilä

## Abstract

**Background:** Recent advances in highly-multiplexed tissue technologies and image analysis tools have enabled a more detailed investigation of the tumor microenvironment (TME) and its spatial features, including tertiary lymphoid structures (TLSs), at single-cell resolution. TLSs play a major part in antitumor immune responses, however, their role in antitumor immunity in ovarian cancer remains largely unexplored.

**Methods:** In this study, we generated a comprehensive single-cell spatial atlas of TLSs in ovarian cancer by extracting spatial topology information from in-situ highly-multiplexed cellular imaging using tissue cyclic immunofluorescence (CyCIF). Our analysis included 44 patients with high-grade serous ovarian cancer (HGSC) from the TOPACIO Phase II clinical trial. We combined spatial and phenotypic features from 302,545 single-cells with histopathology, targeted sequencing-based tumor molecular groups, and Nanostring gene expression data.

**Results:** We find that TLSs are associated with a distinct TME composition and gene expression profile, characterized by elevated levels of the chemokines CCL19, CCL21, and CXCL13 correlating with the number of TLSs in the tumors. Using single-cell feature quantification and spatial mapping, we uncover enriched germinal center (GC) B cell infiltration and selective spatial attraction to follicular helper T and follicular regulatory T cells in the TLSs from chemo-exposed and BRCA1 mutated HGSCs. Importantly, spatial statistics reveal three main groups of cell-to-cell interactions; significantly enriched structural compartments of CD31+ cells, myeloid, and stromal cell types, homotypic cancer cell- and cancer cell to IBA1+ myeloid cell crosstalk, and enriched selective Tfh, Tfr, and Tfc communities with predominant Tfh - GC B cell interactions. Finally, we report spatiotemporal gradients of GC-B cell interactions during TLS maturation, with enriched non-GC B cell attraction towards the GC B cells in early TLSs, and avoidance patterns with selective GC B-cell communities in the TLSs with GCs.

**Conclusions:** Our single-cell multi-omics analyses of TLSs showed evidence of active adaptive immunity with spatial and phenotypic variations among distinct clinical and molecular subtypes of HGSC. Overall, our findings provide new insights into the spatial biology of TLSs and have the potential to improve immunotherapeutic targeting of ovarian cancer.

**What is already known on this topic:** TLSs play a major part in antitumor immune responses, however, their exact role and mechanisms in antitumor immunity are widely unexplored.

**What this study adds:** Our results deepen the understanding of TLS biology including cell-cell interactions and shows how the presence of TLSs is characterized with a distinct TME composition and gene expression profile.

**How this study might affect research, practice or policy:** Overall, our findings provide new insights into the spatial biology of TLSs and have the potential to improve therapeutic options for ovarian cancer.

## Background

Anti-tumor immunity plays a critical role in high-grade serous ovarian cancer (HGSC) therapy responses and clinical outcomes(1,2). Unfortunately, single-agent immune checkpoint blockade therapies in unselected HGSC patient populations have dramatically failed to produce clinical benefits(3). Thus, a better understanding of the tumor microenvironment (TME) is needed to improve immunotherapeutic approaches for ovarian cancer. Interestingly, combined inhibition of Poly-ADP Ribose Polymerase and immune checkpoint inhibition has yielded encouraging results for HGSC patients with an increased proportion of exhausted CD8+ T cells or defective homologous recombination DNA repair(4). Furthermore, the infiltration of CD8+ T cells in *BRCA1/2* mutated (mut) tumors has a prognostic role resulting from their spatial arrangement, highlighting the importance of cell neighborhoods in the TME(5).

Besides T cells, also B cells are found in the TME. Antigen-experienced CD20+ B cells infiltrated together with CD8+ T cells are correlated more strongly to overall survival than CD8+ tumor-infiltrating lymphocytes (TILs) alone in HGSC(6). Infiltrated B cells can together with follicular subtypes of CD4+ T cells form tertiary lymphoid structures (TLSs), which develop in response to prolonged inflammation. TLSs can be diffuse aggregates of lymphocytes, including T cell subsets called follicular T helper (Tfh) and T regulator (Tfr) cells, and maturing B cells, or they can be organized containing B cell follicles and germinal centers (GC) including also highly clonal plasma cells indicating antigen-specific responses.(7,8) In ovarian cancer, tumors with TLSs containing CD8+ TILs and CD20+ B cells are associated with a better prognosis than tumors containing only CD8+ TILs (9). TLSs seem to play a major part in antitumor immune responses, however, their exact role and mechanisms in antitumor immunity are widely unexplored.(10)

To uncover cellular and spatial characteristics of TLSs in ovarian cancer, and to understand TLS shaping factors, we performed highly multiplexed immunofluorescence and single-cell-level image analysis combined with gene expression profiling of tumor samples with recurrent HGSC from the TOPACIO trial(11). Altogether our study reveals distinct gene expression programs, cellular compositions, and spatial cell-cell interactions in HGSC associated with TLS maturation and clinical and molecular patient groups.

## Materials and methods

### Study cohort

The study consisted of 44 patients participating in the TOPACIO Phase II clincal trial (4)(11). Formalin-fixed paraffin-embedded (FFPE) tumor samples were collected either at the time of diagnosis (chemo-naïve) or after neoadjuvant platinum-based chemotherapeutics (chemo-exposed). The H&E stains were available on 42/44 tumor samples (4). A trained pathologist (A.L.) annotated the tumors for TLSs from the H&E digital sections, and the TLSs were further labeled into three categories based on their location (intratumoral, peritumoral, or “both”) to the invasive tumor margin. TLSs were detected in 26/44 tumor samples. A subcohort of 19/44 tumor samples was analyzed with single-cell image analysis using CyCIF, in which ten tumor samples contained TLSs and nine did not. The TLS status obtained from H&E stains differed in 3/16 samples obtained from the CyCIF analysis,due to that the H&E and CyCIF assays were not performed from consecutive tissue sections. In the CyCIF subcohort, three samples were excluded due to suboptimal tissue quality. Characteristics of the clinical data and the TLS status are shown in **Table S1**.

### Cyclic immunofluorescence

5µm FFPE sections were stained using highly multiplexed cyclic immunofluorescence (CyCIF)(12)(4). Briefly, the FFPE tissue sections were cyclically stained with antibodies (**Table S1**) and scanned with the RareCyte CyteFinder scanner. Scanned image files were corrected using BaSiC and stitched and registered using the ASHLAR algorithm(13). The single-cell proportions of the WSIs were utilized from our previous study(4).

### Single-cell image analysis of TLSs

TLS aggregates were cropped from the whole-slide images (WSIs) using ImageJ(14) utilizing morphology and CD20 signal for TLSs. The nuclei were segmented using StarDist’s versatile (fluorescent nuclei) model(15). The median fluorescence intensity (MFI) for each cell mask was computed using the Bio-Formats library in Matlab(16). To filter out cells with diminishing signals during the CyCIF protocol, we used the repeated DNA MFI measurement. The possible lost cells were visualized in Napari(17). **Table S1** shows the number of images analyzed including the quality assessment of each TLS.

The quantified single-cell data were normalized, phenotyped, and further analyzed using Scimap(18) (https://scimap.xyz/). For the markers used for phenotyping, the positive thresholds were set manually after which the signal intensities of the cropped TLSs were normalized per tumor sample. For the markers not used in the cell phenotyping, a Gaussian mixture model was used to normalize the signal intensities. **Table S2** summarizes the manual thresholds and shows the workflow of the cell phenotyping. In total, 302,545 single cells were identified and phenotyped based on their marker positivity. A minority of double negative (CD3+CD4-CD8-) T cells (334 cells, 0.11%) and granulocytes (CD15+, 114 cells, 0.05%) were removed from further analysis due to low cell counts. The schematic pipeline of the image analysis is shown in **Figure 1A**.

**Figure 1:**
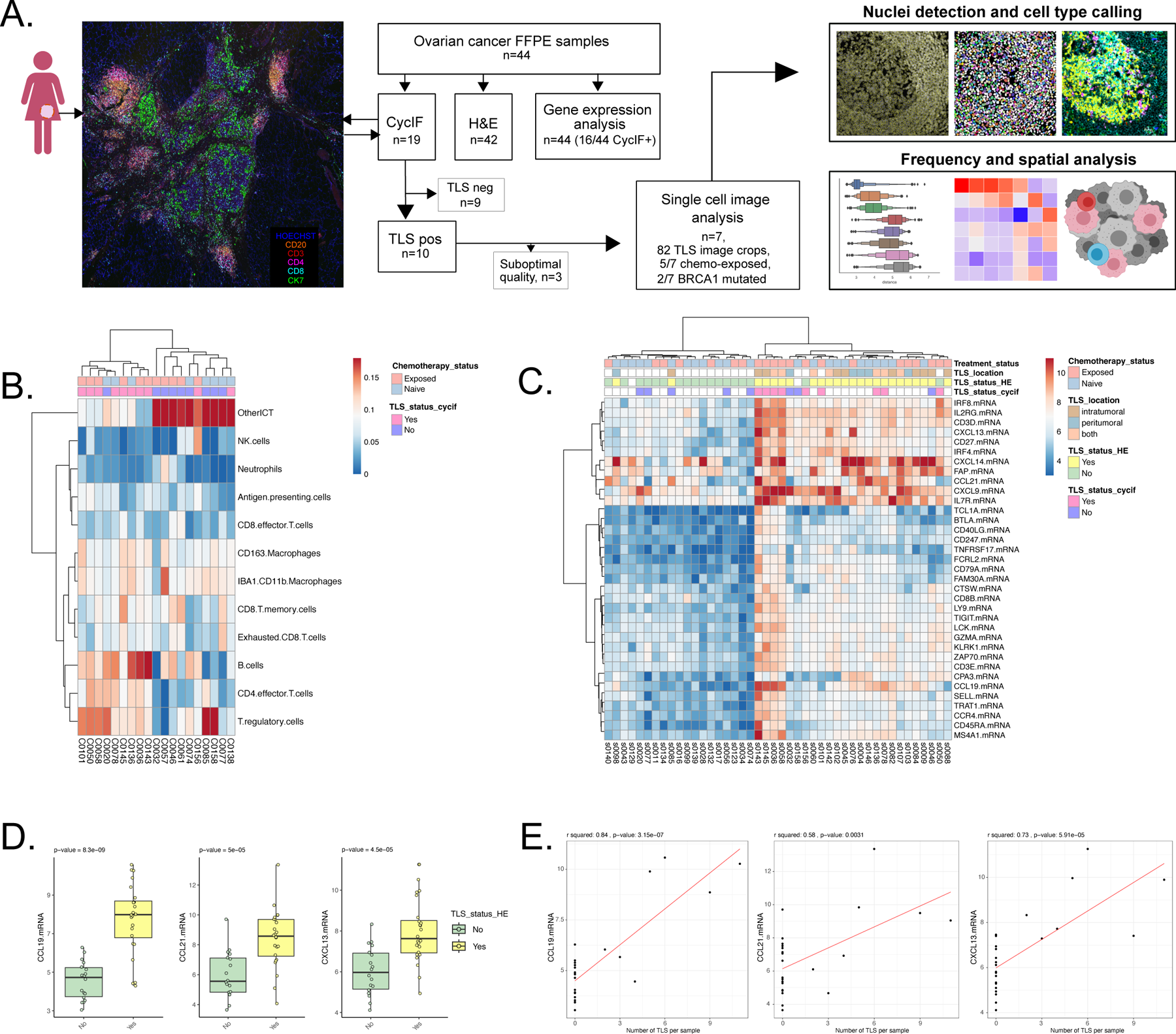
The presence of TLSs is characterized by distinct TME and gene expression profiles. (A) Schematics demonstrate the study design. The inclusion of tumor samples with TLSs for single-cell image analysis is shown in the flow diagram. (B) Cluster heatmap displaying correlation of TLS status with CyCIF WSI cell type proportions obtained from Färkkilä et al (4). The color in the box corresponds to the proportion of cell type in a given tumor. Heatmap clustering was performed with Ward D2 linkage. (C) Cluster heatmap of gene expression analysis of 44 tumors using a p-value cutoff of −log10 of 3.0 and a log2FC of >1.5 or <-1.5 correlates by the TLS status. The color in the box corresponds to the mRNA level in a given tumor. Heatmap clustering was performed with Ward D2 linkage. Genes differentially expressed within the treatment comparison were filtered out. (D) Box and whisker plot of *CCL19*, *CCL21*, and *CXCL13* mRNA level by the TLS status obtained from digitized H&E stained digital section of HGSC. The P-value was calculated with an unpaired Wilcoxon test. (E) Pearson correlation test of *CCL19* and *CXCL13* mRNA level against TLS number in CyCIF annotated tumors (*CCL19*, p=8.3e-09; *CCL21*, p=5e-05; CXCL13, p=4.5e-05). Sample 0078 contained 48 TLSs and was excluded from the correlation tests as an outlier.

### Single-cell spatial analysis

The neighborhood analyses were adopted from Scimap using Jupyter Notebook with Python. First, the average shortest spatial distances between the different cell types were calculated to create a network of cellular distances across the TLSs. Secondly, the K-nearest neighbors’ method with 10 nearest cells was applied, to determine the cell-type-specific cellular communities for every cell and to identify the spatial patterns of attraction and avoidance between the given cell types(18).

### Gene expression profiling and pathway analysis

The Nanostring gene expression profiling using a PanCancer IO 360 Gene Expression Panel plus 30 DNA repair genes was used in a cohort of 44 patients on FFPE tumor sections obtained from the same FFPE blocks which were used in the CyCIF experiment(4). The data were normalized to 20 housekeeping genes and analyzed through the NanoString NSolver Advanced Analysis Platform, including pathway enrichment scores. An unpaired t-test was used for the differential gene expression analysis using a p-value cutoff of −log10 of 3.0 and a log2FC of >1.5 or <-1.5. Genes affected by the treatment status were filtered out (a cutoff of −log10 p-value of 3.0 and a log2FC of >2.5 or <−2.5). Differentially expressed KEGG pathways were detected using a −log10 p-value of 2.5 and a log2FC cutoff of >2.5 or <−2.5.

### Statistics

Either an unpaired Wilcoxon or a t-test was used to compare two different categories. P-values of <0.05 were considered statistically significant. For the RNAseq analysis, the distribution of the data was approximately normal based on the Shapiro-Wilk test and an unpaired T-test was applied. A Pearson correlation test was used for certain differentially expressed genes. Euclidean dissimilarity distances and Ward’s agglomerative clustering were used in the correlation plots. P-values for the cell-type-specific cellular communities were either calculated by subtracting the permuted mean from the observed mean divided by the number of permutations described by Schapiro et al(19) or by using an unpaired Wilcoxon test to compare the cell-cell spatial interaction of given cell types between two categorical groups.

## Results

### The presence of TLSs is reflected in the overall TME cell composition in HGSC

Using highly-multiplexed imaging, we identified infiltrations of B cells (CD20+) colocalizing with CD3+CD4+ T helper cells forming TLSs in 10 out of 19 (53%) HGSC samples (**Figure 1A**). First, we compared how the presence of TLSs was reflected in the proportions of different immune cell subtypes in the whole slide image (WSI) CyCIF data. Agglomerative hierarchical clustering resulted in two larger clusters that followed the presence of TLSs (**Figure 1B**). Ovarian cancers with TLSs had a higher proportion of CD20+ B cells (p=0.016) and CD4+ T helper cells (CD3+CD4+FOXP3-, p=0.010) as compared to tumors without TLSs (**Figure S1A**).

### The presence of TLSs is characterized by a distinct gene expression profile of CCL19, CCL21, and CXCL13

We next tested whether the presence of TLSs was associated with gene expression profiles of the tumors using expression data of 800 genes from the Nanostring platform from 44 patients. Based on histopathology, TLSs were identified in 24 tumors whereas in 18 tumors they were absent (**Figure S1B)**. Treatment with chemotherapy has been associated with an increase in TLS formation in various cancer types(10,20), however here the presence of TLSs did not correlate with the chemotherapy status (12/21 chemo-naïve vs 12/21 chemo-exposed), *BRCA1/2* mutation status (4/8 *BRCA1/2*mut vs 20/34 *BRCA1/2* wild-type), or the outcomes of patients (11/17 clinical benefit vs 13/25 without benefit) in the TOPACIO trial.

Altogether, we identified 36 genes upregulated in tumors with TLSs, which remained after filtering for chemo-exposure status. (**Figure S1C)**. Again, the tumors clustered into two major clusters correlating with the presence of TLSs in histopathological assessment (**Figure 1C**), while chemotherapy exposure was not associated with the main clusters. No clear gene expression patterns were observed based on TLS location (intratumoral, peritumoral, or “both”) (**Figure 1C**). Interestingly, we observed a significant upregulation of several genes encoding for chemoattractants responsible for lymphocyte migration into TLSs (*CCL19*, p=8.3e-09; *CCL21*, p=5e-05; CXCL13, p=4.5e-05; **Figure 1D**). Furthermore, the expression levels of *CCL19* and *CXCL13* also significantly correlated with the number of TLSs in the tumors (*CCL19*, R-squared=0.84, p=3.2e-07; *CXCL13*, R-squared=0.73, p=5.9e-05; **Figure 1E**).

We next explored the gene expression patterns using KEGG pathways of the differentially expressed genes. We observed that the tumors with TLSs showed increased activity in immunoregulatory interactions between lymphoid and non-lymphoid cells, TCR signaling, IL2, 4, and 10 signaling, and costimulation by the CD28 family. Again, the tumors clustered according to the TLS status, implicating distinct gene expression programs in tumors with TLSs (**Figure S1D**).

### CyCIF reveals heterogeneity of TLS composition at single-cell resolution

We next set out to characterize the composition of the TLSs at single-cell resolution. In total, 302,545 single cells from 82 TLSs from WSIs encompassing 7 different HGSC patients were subjected to further single-cell analyses (**Figure 1A, Table S1**). We phenotyped the cells to different subsets of CD4+ T helper cells, regulatory T cells, CD8+ T killer cells, CD20+ B cells, myeloid, stromal, endothelial, and cancer cells, with a specific focus on follicular T cells by utilizing probability-based marker expressions (**Figure 2A, Table S2**). Consistent with the TLS biology, B cells and CD4+ T cells encompassed over 50% of the cells identified in the TLSs, followed by myeloid, CD8+ T, and stromal cells (**Figure 2B**). In unsupervised hierarchical clustering, the TLSs formed four main clusters enriched either in T-cell populations, cancer and myeloid cells, B-cells, or stromal cells. Interestingly the individual TLSs showed marked heterogeneity in the main cell lineage composition between the patients (**Figure 2C**). Notably, the samples with the largest frequencies of B cells did not possess the largest frequencies of T cell subsets and had relatively low frequencies of other cells, consistent with the WSI data (**Figure 1B,2C**).

**Figure 2:**
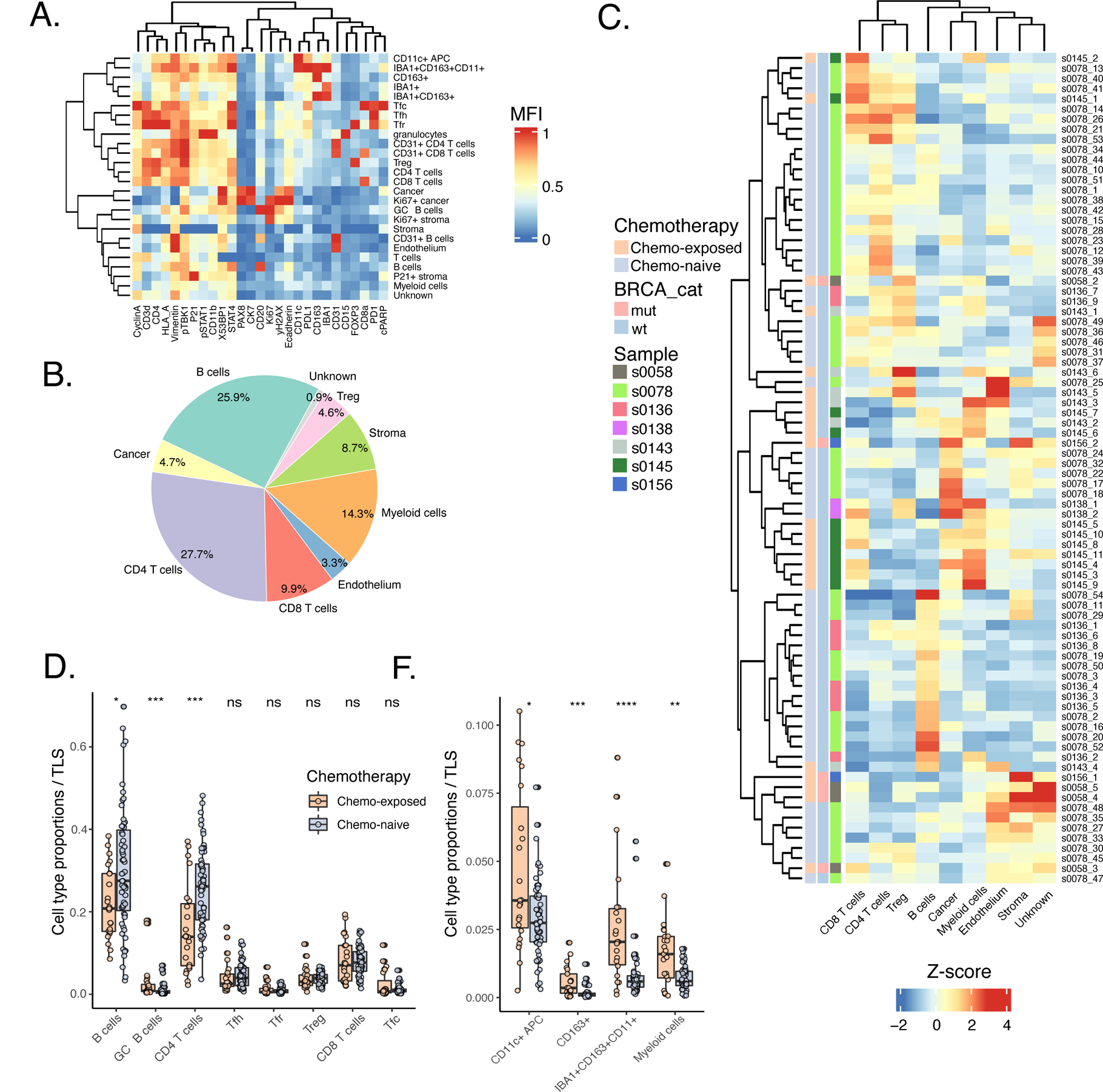
Cell composition of TLSs and increased proportion of GC B cells in chemo-exposed tumors. (A) Heatmap showing scaled mean fluorescence intensities of different markers detected from CyCIF images per cell type in TLSs after probability-based phenotyping. (B) Proportions of main cell lineages of all cells (%) in TLSs from CyCIF images are shown using a pie chart. (C) Cluster heatmap of proprotions of main cell lineages in individual TLSs shows heterogeneity between individual TLSs and samples. Clustering was achieved using the complete method. Scaling to z-scores was performed cell lineage-specifically. (D) Lymphocyte compartment in TLSs from chemo-exposed tumors shows an increased proportion of GC B (CD20+Ki67+, p=8e-04) cells and naïve T helper (Th, CD3+CD4+PD1-FOXP3-) cells (p=4e-04) whereas the proportion of non-GC B (CD20+Ki67-, p=0.02) cells was decreased compared to chemo-naïve TLSs (unpaired Wilcoxon test). No statistical differences were observed in proportions of follicular T cells (Tfh, CD3+CD4+PD1+FOXP3-; Tfr, CD3+CD4+PD1+FOXP3+; Tfc, CD3+CD8+PD1+) and cytotoxic T (Tc, CD3+CD8+PD1-) cells between the subtypes. (E) chemo-exposed TLSs had a higher proportion of all myeloid subsets compared to chemo-naïve TLSs (CD11c+ APCs, p=0.02; CD163+ macrophages, p=7e-04; IBA1+CD163+CD11c+, p=1e-05; unspecified myeloid cells, p=0.002. In (C) and (D), p-values were calculated with an unpaired Wilcoxon test.

### Chemotherapy-exposed TLSs are enriched in germinal center B cells

Even though there was no difference in TLS prevalence between chemo-exposed and naïve tumors, we hypothesized that chemotherapy would influence the TLS composition. First, we compared the proportions of the main cell lineages within the TLSs and noticed that the chemo-exposed TLSs seem to have a reduction of overall B cells (CD20+; p=0.01) and CD4 T helper cells (CD3+CD4+FOXP3-; p=1e-04), and enrichment of endothelial cells (CD31+; p=0.01) and myeloid cells (p=2e-09) as compared with the chemo-naïve TLSs (**Figure S2**). A more detailed analysis of the lymphocyte compartment revealed that the chemo-exposed TLSs had a significantly higher proportion of proliferative GC-related CD20+Ki67+ B cells (GC B; p=8e-04 and subsequently the fraction of non-GC (CD20+Ki67-) B cells (p=0.02; **Figure 2D**) was reduced. The proportion of CD3+CD4+PD1-FOXP3-(CD4 T) cells was lower in the chemo-exposed TLSs (p=4e-04). The other T cell subsets including PD1+ populations did not differ between the subgroups. Traditionally, follicular T helper (Tfh) and regulator (Tfr) cells are <characterized by chemokine receptor CXCR5 in addition to canonical T cell markers, however, their spatial localization in the TLS follicles, as shown later in **Figure 3B (i-iii, vi)**, makes it plausible to annotate the CD3+CD4+PD1+ as Tfh and CD3+CD4+PD1+FOXP3+ as Tfr cells. In the TLS follicle, the PD1+ cells were Tfh cells and not GC B cells (**Figure 3B iii-iv**).

**Figure 3:**
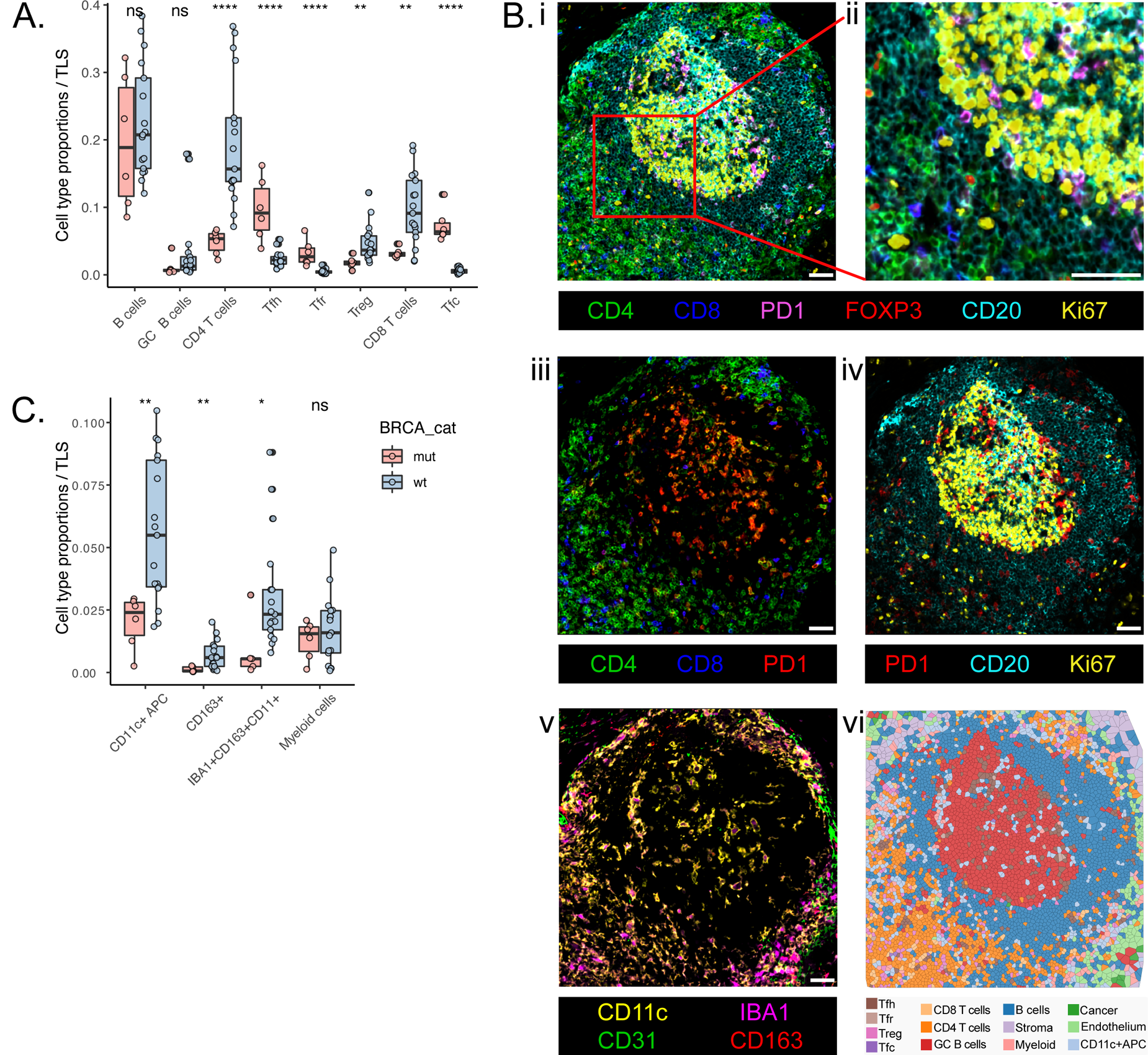
The proportion of follicular T cells is increased in *BRCA1* mutated TLSs. (A) The proportion of lymphocytes in chemo-exposed TLSs shows the increased proportion of Tfh (p=8e-05), Tfr (p=5e-05), and Tfc cells (p=2e-05) cells and the decreased proportion of Th (p=2e-05), Treg (p=0.002), and Tc (p=0.002) cells in *BRCA1* mutated tumors compared to wild-type (an unpaired Wilcoxon test). No statistical difference in B cell populations was observed. (B) CyCIF images with scale bars of 50 µm demonstrate the organization of a TLS and the location of its cell types. i) An example of TLS with GC B (CD20+Ki67+) cells containing follicle showing how follicular (PD1+, pink) T cells are located mostly inside or vicinity of the follicle. ii) A magnification showing the interface of GC and mantle zone. iii) PD1+ (red) cells are mostly CD4+ (green) Tfh cells (orange) or CD8+ (blue) Tfc cells (purple). iv) Ki67+ (yellow) cells are mostly CD20+ (cyan) GC B cells found in the follicle. v) CD11c+ (yellow), IBA1+ (pink), and CD163+ (red) myeloid cell types and CD31+ endothelial cells are seen mostly outside the follicle. Some CD11c+ APCs are found in the follicle. vi) A Voronoi plot using segmented cells shows typical locations of different cell types. For simplicity, different myeloid cell types were pooled as “Myeloid”, cancer cells as “Cancer”, and stromal cells as “Stroma”. CD31-positive CD4 T, CD8 T, and B cells were included in “CD4 T”, “CD8 T cells”, and “B cells”, respectively. (C) All myeloid cell subsets were decreased in chemo-exposed *BRCA1* mutated TLS compared to chemo-exposed wild-type TLSs (CD11c+ APCs, p=0.003; CD163+, p=0.006; IBA1+CD163+CD11c+, p=0.01). P-values were calculated with an unpaired Wilcoxon test.

The myeloid cells (CD11b+/CD11c+/CD163+/IBA1+) were further divided into CD11c+ antigen-presenting cells (APC), CD163+ and IBA1+CD163+CD11c+ macrophages, and unspecified myeloid cells (**Table S2**). We observed that the chemo-exposed TLSs had an enriched proportion of all myeloid subsets as compared to the chemo-naïve TLSs (CD11c+ APCs, p=0.02; CD163+ macrophages, p=7e-04; IBA1+CD163+CD11c+, p=1e-05; unspecified myeloid cells, p = 0.002; **Figure 2E**).

### Enriched infiltration of follicular T-cells in the TLSs of *BRCA1* mutated tumors

To enhance our understanding of the TME after chemotherapy, we focused on the TLSs from chemotherapy-exposed tumors across different molecular subgroups of HGSC. Previous studies have shown that mismatch repair-deficient colorectal cancers have an increased prevalence of TLSs with more mature phenotypes(20), but this was not observed in *BRCA1*mut breast cancer patients(21). *BRCA1* mutation could nevertheless alter TLSs on a cellular level, which led us to compare TLS cell composition of chemotherapy-exposed tumors between *BRCA1*mut TLSs to TLSs from tumors with no alterations in homologous recombination pathways (wild-type).

TLSs from *BRCA1*mut had an increased infiltration of Tfh (p=8e-05) and Tfr cells (p=5e-05), while unspecified CD4 T cells (p=2e-05) and CD3+CD4+FOXP3+PD1-regulatory T cells (Tregs; p=0.002) were decreased compared to wild-type TLSs (**Figure 3A**). Interestingly, we observed divergent CD3+CD8+ T cell infiltration patterns in the TLSs of *BRCA1*mut and wild-type tumors. The TLSs in *BRCA1*mut tumors showed an enriched infiltration of PD1+CD8 T cells (p=2e-05; **Figure 3A**), which are likely follicular cytotoxic T (Tfc) cells since they spatially located inside or near the TLS follicle (**Figure 3B iii, vi**), while the fraction of PD1-negative CD8 T cells was reduced (p=0.002) compared to wild-type TLSs consistent with general observations from the tumor-infiltrating lymphocytes(22).

Earlier we reported reduced infiltration of CD11c+ APCs and IBA1+CD163+CD11c+ macrophages in the *BRCA1*mut tumors as compared to wild-type(5). Similarly in the TLSs, we observed lower fractions of both CD11c+ APCs (p=0.003), CD163+ (p=0.006) and IBA1+CD163+CD11c+ (p=0.01) macrophages in the *BRCA1*mut TLSs (**Figure 3C**). The reduced infiltration of myeloid cells in the TLSs could also rise from highly expanded Tf cells reported above because no changes in different myeloid cell subpopulations were seen within the myeloid compartment alone (**Figure S3**). Consistently, the myeloid cells were mostly located outside of the TLS follicle, yet the CD11c+ APCs were also found inside the GC B cell-rich follicle (**Figure 3B v-vi**).

### Selective spatial interactions of follicular T cells and GC B cells

To investigate the cell-to-cell spatial interactions within the TLSs, we next calculated the average shortest spatial distances between different cell types to create a network of cellular distances. We found that cells of the same type were more likely to cluster together, indicating spatial clustering of the cells of the same type towards each other (**Figure S4A**). Based on average distances between cell types, the cellular interactions were clustered into three main categories of cellular networks: 1) homotypic interactions (i.e., cells adjacent only with the same cell type, and distant from other cell types), 2) heterotypic interactions (i.e. cells closely interacting with many other cell types), and 3) selective interactions (i.e. cells adjacent mostly with the same cell types, but also with a subset of other cells) (**Figure 4A**). The homotypic interactions involved cancer cells (CK7+ or PAX8+), proliferative (Ki67+) cancer and stromal cells, CD163+ myeloid cells, and CD31+ lymphocytes (CD31+ CD4 T cells, CD8 T cells, or B cells) which may represent trafficking lymphocytes (23) since they were mostly found around CD31+ endothelial cells including HEVs confirmed by visual inspection and their cellular communities (as shown later in **Figure 4C,5C**). The heterotypic interactions consisted of cell types that closely interacted with a variety of different cell types across TLSs and contained non-GC B cells, unspecified CD4 and CD8 T cells, Tregs, CD11c+ APCs, stromal, and endothelial cells. Selective cellular interactions within the TLSs consisted of various myeloid cell types (IBA1+CD163+CD11c+, IBA1+CD163+, unspecified myeloid cells), GC B cells, and follicular T cells (Tfh, Tfr, and Tfc cells). Interestingly, the spatial interactions of the GC B and follicular T cells formed an independent subcluster. Consistently, the Tfh cells were the only cell type adjacent and interacting with the GC B and non-GC B cells within the selective interactions network. The spatial projection of the distinct cell-cell interactions on the underlying highly-multiplexed image confirms the presence of the distinct spatial interaction networks in the TLSs (**Figure 4B**).

**Figure 4:**
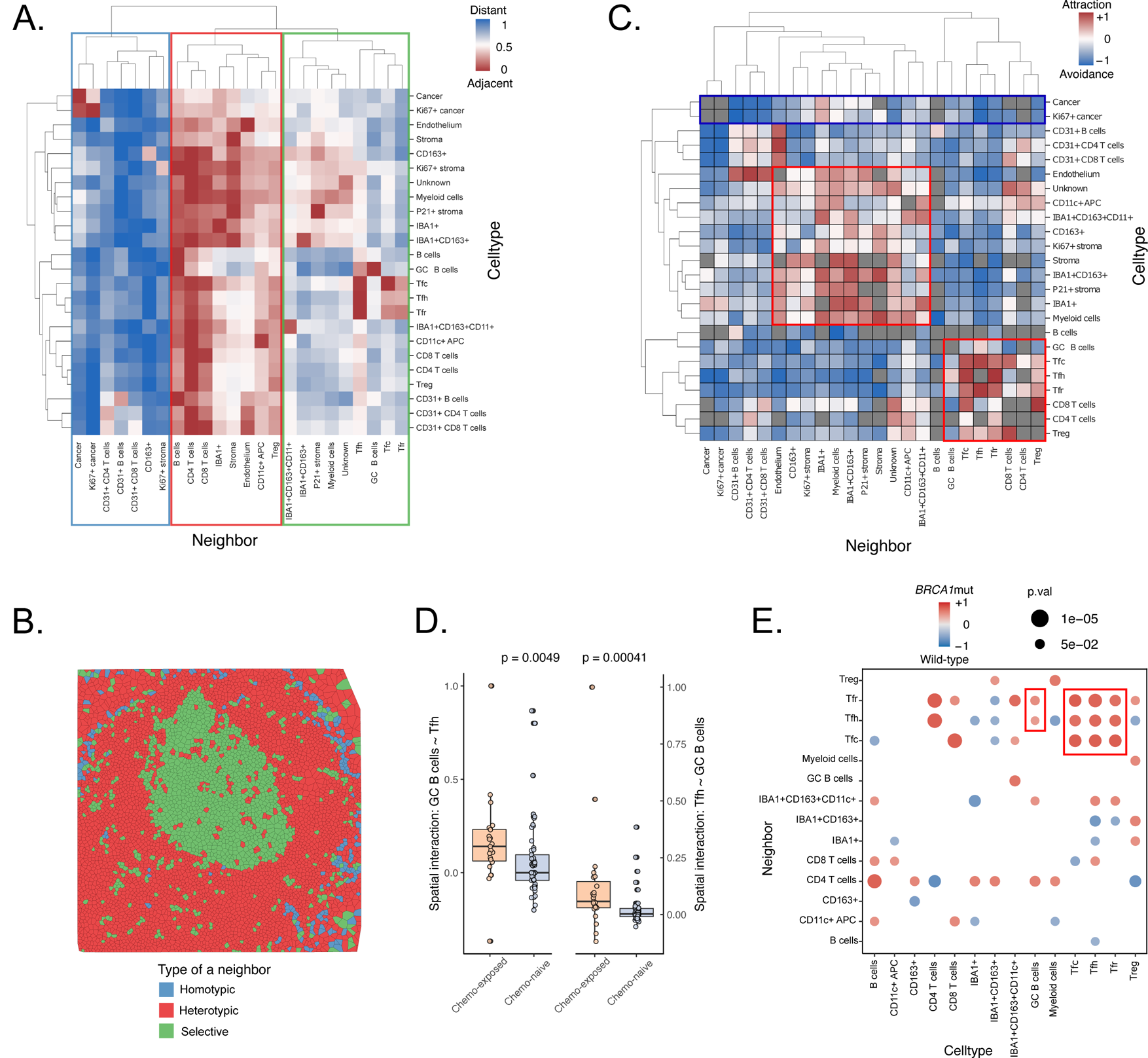
TLSs harbor unique cell type-specific neighborhoods. (A) The average shortest distances between different cell types are presented as a heatmap. Cell types were clustered into three different interaction networks: 1) homotypic cells, i.e., cells adjacent only with themselves (blue), 2) homotypic cells, i.e., cells interacting with most cell types (red), and 3) selective cells, i.e., cells interacting with certain cell types (orange). Cell-cell distances are in log scale, where the red color means that cell types are adjacent whereas the blue color means they are distant. (B) A Voronoi plot of the TLS displayed in Figure 3B illustrates how separate interaction networks are located in a TLS. Each cell was annotated based on the interaction network identified in **A**. (C) The immediate cell-type-specific cellular communities of the TLS cell types show restricted communities for i) lymphocytes and ii) myeloid, stromal, and endothelial cells (highlighted with red rectangles). The red color reflects attraction between given cell types, whereas the blue color means avoidance compared to random. P-values were calculated by subtracting the permuted mean from the observed mean divided by the number of permutations. (D) Chemo-exposed TLSs harbor more cell-cell interactions between GC B cells and Tfh cells than chemo-naïve TLSs, both when Tfh cells neighbor GC B cells (left, p=0.005), and when GC B cells neighbor Tfh cells (right, p=0.0004). The spatial interaction between the cell types was compared using an unpaired Wilcoxon test. (E) A dot plot summarizes cell-type-specific immediate cellular communities by comparing the cell-cell spatial interactions between the *BRCA1*mut and wild-type TLSs with an unpaired Wilcoxon test. GC B cells have more spatial interactions with Tfh, Tfr, and Tfc cells in TLSs from tumors with *BRCA1*mut than wild-type. The size of the dot reflects statistical significance. In **C-E**, the 10 nearest neighbors were used to determine the immediate communities for every cell.

**Figure 5:**
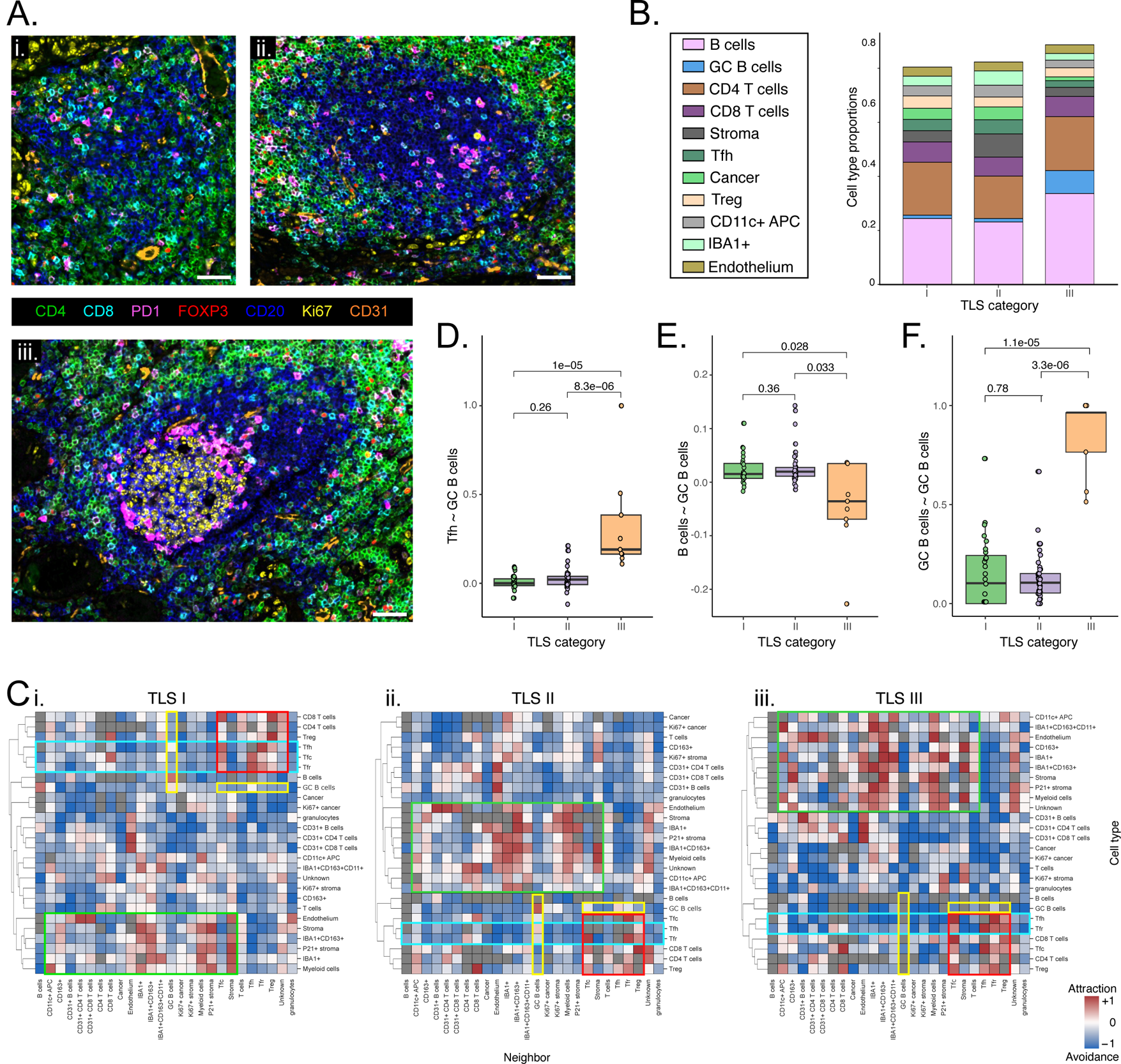
Cell composition and spatial interactions change during the development of TLS. (A) CyCIF images with scale bars of 50 µm demonstrate structural characteristics of each TLS category: i) TLS-I: unorganized aggregates of B (CD20, blue), CD4 (green), and CD8 (cyan) T cells; ii) TLS-II: larger aggregates of lymphocytes with a B cell follicle in the middle. iii) TLS-III: an organized TLS with a B cell follicle containing a Ki67-positive (yellow) GC infiltrated with the PD1+ (pink) Tfh cells. (B) A stacked bar plot showing the proportions of different cell types of all cells. The only statistical changes in cell type proportions between TLS categories were observed in the Ki67+ cancer and GC B cells as shown in more detail in Figure S5A-B. The CD31+ lymphocytes and certain stromal, myeloid, and cancer cell subsets are not shown due to their low proportions. (C) The immediate cell-type-specific cellular communities for separate TLS categories are summarised using cluster heatmaps. P-values were calculated by subtracting the permuted mean from the observed mean divided by the number of permutations. The red color reflects an attraction between given cell types, whereas the blue color means avoidance compared to random. The Tfh, Tfr, and Tfc cells clustered together with the unspecified CD4 and CD8 T cells, and Tregs throughout the TLS maturation phases (the red rectangles), however, the avoidance of Tfh and Tfr cells towards other cells than lymphocytes got more prominent during the maturation from lymphocyte aggregates (TLS-I) towards the TLSs with GCs (TLS-III, the blue rectangles). The spatial communities involving the GC B cells contained only few significant attractions (the yellow rectangles). The myeloid and stromal cells where interacting mostly with themselves (the green rectangles). (D) The cellular communities of the Tfh cells had increased interactions with the GC B cells in the TLSs with GC formation as compared to the TLSs without GCs (TLS-I vs TLS-III, p=1e-05; TLS-II vs TLS-III, p=8.3e-06). (E) The cellular communities of the non-GC B cells had less interactions to the GC B cells in TLSs with GC formation as compared to the TLSs without GC (TLS-I, p=0.028; TLS-II, p=0.033). Within the GC B cells communities, the GC B cells were interacting more with themselves in the TLSs with GC as compared to TLSs without GC (TLS-I, p=1.1e-05; TLS-II, p=3.3e-06). In **D-F**, the spatial interactions between the cell types were compared using an unpaired Wilcoxon test. The 10 nearest cells were used in **C-F** to define immediate communities for every cell.

Next, we investigated the cell-type-specific cellular communities of 10 spatially closest neighbors for every cell. To reveal patterns of attraction or avoidance, we computed a statistical measure of how likely a given cell type is to be found in the proximity of another cell type as compared to random(19). We identified distinct significantly enriched cellular communities which clustered into three groups (**Figure 4C**). First, we observed a significant attraction of cancer cells towards each other, consistent with observations in **Figure 4A**. The only statistically significant neighbor cell type for cancer cells in TLSs were IBA1+ myeloid cells supporting the role of myeloid cells in regulating anti-tumor immunity in HGSC. Secondly, we observed significantly enriched communities of CD31+ cells, myeloid, and stromal cell types (**Figure 4C**). We found that the CD31+ endothelial cells including HEVs, together with the myeloid and stromal cell types showed significant attraction and co-occurrence with potential structural functions for the TLSs, consistent with the stromal cells and HEVs being especially critical for TLS development and the migration of lymphocytes to the TLS site(24). The myeloid, endothelial, and stromal cells were also mostly found in statistically significant spatial avoidance of the GC B cells, outside of the follicle supporting their function in the TLS structures (**Figure 4C**), as also visible in **Figure 3B v-vi**. The third cluster of significant attractions was formed by lymphocytes, highlighting the critical role of spatial lymphocytic interactions in TLS function. Interestingly, the Tfh, Tfr, and Tfc cells formed communities of significant spatial attraction, whereas the cellular neighborhoods of unspecified CD4 and CD8 T cells, and Tregs were more scattered in the TLSs. Of note, the Tfc cells were significantly enriched in the T cell neighborhoods but showed avoidance patterns towards cancer and myeloid cells supporting the notion that the CD3+CD8+PD1+ cells in TLSs here represent a follicular subset of CD8 T (Tfc) cells rather than the classical exhausted CD8 T cells. Importantly, the Tfh cells were the only cells forming statistically significant attractions to the GC B cells (**Figure 4C**) highlighting the importance of selective spatial crosstalk between the GC B and Tfh cells for the GC reaction.

### Enriched spatial interactions of follicular T cells and GC B cells in chemo-exposed and *BRCA1*mut HGSCs

Since the cell composition of TLSs differed between chemo-exposed and naïve tumors (**Figure 2C-D**), we next investigated whether the cell-type-specific cellular communities presented in **Figure 4C** were affected by the chemotherapy status. We observed drastic differences in immediate cell-cell interactions especially for the GC B and Tfh cells. The cell communities of the GC B cells included increased attractions to the Tfh (p=0.005, **Figure 4D**) and Tfr cells (p=0.004, **Figure S4B**) in the chemo-exposed TLS as compared to the chemo-naïve TLSs. Similarly, the cell communities of the Tfh cells harbored increased attractions to the GC B cells (p=0.0004, **Figure 4D**) and to the non-GC B cells (p=0.019, **Figure S4B**) in the chemo-exposed TLS as compared to the chemo-naïve TLSs. Even though the myeloid cell subsets were increased in the chemo-exposed tumors, we did not detect evident patterns of changes in the myeloid communities between TLSs of different chemotherapy statuses (**Figure S4C**).

We then focused the analysis to the chemo-exposed TLSs and compared the cell-type-specific cellular communities (knn=10 cells) of *BRCA1*mut and wild-type TLSs. We observed that, in addition to the enrichment of follicular T cell populations (**Figure 3A**), the *BRCA1*mut TLSs also had increased spatial attractions between the GC B and all follicular T cell populations as compared to the wild-type TLSs (**Figure S4D**). Of note, in the wild-type TLSs, the GC B cells had significant attractions only with the Tfh cells. The spatial attraction between GC B cells and Tfh/Tfr cells was found to be significantly increased in *BRCA1*mut tumors compared to wild type (p=0.04 and p=0.02, respectively, as shown in **Figure 4E**). Further, the cellular communities of the Tfh, Tfr, and Tfc cells had increased attraction with each other in TLSs from the *BRCA1*mut tumors as compared to the wild-type (**Figure 4E**). However, no significant difference was observed in the interactions between GC B cells and Tfc cells.

### Spatial interaction dynamics during TLS maturation

To shed light on the dynamic changes in the development of TLSs, we classified the TLSs into three categories using morphological characteristics(25). Over one-third of all TLSs consisted of small aggregates of lymphocytes without an evident B cell follicle (TLS-I, 34.1%, 28/82, **Figure 5A i**). TLSs with larger aggregates of lymphocytes containing a B cell follicle without GC formation consisted of over half of all TLSs (TLS-II, 54.9%, 45/82, **Figure 5A ii**), whereas a GC, confirmed with Ki67-positivity, was seen in 11.0% of the TLSs (TLS-III, 9/82, **Figure 5A iii**). Although morphological changes were observed among TLSs, the only significant differences in cell compositions between TLS categories were found in Ki67+ cancer cells and GC B cells (**Figure 5B**). Specifically, the proportion of Ki67+ cancer cells was significantly higher in TLS-I than in TLS-II (p=0.006) and TLS-III with GCs (p=0.003), while no statistical differences in cell composition were observed between TLS-II and TLS-III (**Figure S5A**). In line with TLS maturation, TLSs with GC formation (TLS-III) showed enrichment of GC B cells as compared to TLS aggregates without a B cell follicle (TLS-I, p=0.003) or TLSs with a B cell follicle but no GC (TLS-II, p=0.003, **Figure S5B**).

Next, we used spatial statistics to examine the spatial dynamics of cell-type-specific cellular communities (knn=10 cells) during TLS maturation (Figure 5C-F). The Tfh, Tfr, and Tfc cells formed a large spatial cluster with CD4 and CD8 T cells, and Tregs throughout all TLS maturation phases (**Figure 5C**). However, Tfh and Tfr cells showed increasing avoidance of other cell types during TLS maturation, indicating the segregation of the follicular T-cell communities in TLS-III. Both GC and non-GC B cells formed a separate spatial cluster next to the T cell cluster in all TLS categories. In TLS-IIs, the GC B cells attracted Tfh cells, while in TLS-Is, they only attracted non-GC B cells in addition to themselves, supporting the development of B cell follicles. The cellular communities of different myeloid cell types primarily consisted of other myeloid cell subpopulations, which mostly showed avoidance patterns towards lymphocyte populations. In the more mature TLSs (TLS-II and TLS-III), stromal and endothelial cells also accompanied the myeloid cell communities (**Figure 5C**), supporting the structural maturation of the TLSs.

The categories of TLSs were not solely associated with cell compositions (**Figure S5C**), highlighting the importance of cell-type-specific cellular communities and cell-cell interactions in TLS development (**Figure 5C**). Since GC B cells exhibited only a few statistically significant attractions to other cell types, we investigated whether cell-type-specific cellular communities involved more interactions with GC B cells in specific TLS categories. Tfh cells had significantly enriched interactions with GC B cells in TLSs with GC formation compared to those without (TLS-I, p=1e-05; TLS-II, p=8.3e-06, **Figure 5D**). No difference in interactions between Tfh and GC B cells was observed between TLS-I and TLS-II, even though a B cell follicle was already present in the TLS-II. Similarly, GC B cell communities showed a trend of enriched attraction to Tfh cells in TLS-III (**Figure S5D**). Other follicular T cells (i.e., Tfr and Tfc cells) were not truly attracting or avoiding GC B cells in any TLS category, although Tfc cell communities in the TLS-IIs had slightly more interactions with GC B cells than in TLS-IIIs (p=0.017, **Figure S5E-F**). Interestingly, the cellular communities of non-GC B cells had fewer interactions with GC B cells in TLSs with GC formation than those without (TLS-I, p=0.028; TLS-II, p=0.033, **Figure 5C,E**). Conversely, the GC B cells had significantly increased selective interactions with themselves in TLSs with GC (TLS-III) compared to those without (TLS-I, p=1.1e-05; TLS-II, p=3.3e-06, Figure 5F), while spatial interactions were similar in TLS-I and TLS-II, highlighting the unique spatiotemporal dynamics of the GC reaction during TLS maturation.

## Discussion

The recent developments in the highly-multiplexed tissue technologies and image analysis tools have enabled a more detailed investigation of the TME and its spatial structures such as TLSs at single-cell resolution. Here, using single-cell feature quantification and spatial statistics in high-plex marker space, we present a detailed map of TLSs in ovarian cancer. We show that spatially and functionally heterogeneous TLSs are present in HGSCs regardless of the chemotherapy exposure. Further, we revealed that TLS structures were associated with a distinct TME composition and tumor gene expression signature with elevated expression of chemokines CCL19, CCL21, and CXCL13. Spatial and structural analysis revealed unique cellular interactions and dynamics during TLS maturation in the ovarian cancer TME.

By combining single-cell profiling of TLS, whole-slide imaging (WSI), and gene-expression profiling, we explored the cellular composition and gene expression signatures indicative of active adaptive anti-tumor immunity. We observed that the presence of TLSs correlated with an increased proportion of B cells and T helper (Th) and regulator (T reg) cells in whole slides of tumor samples obtained with CyCIF(4). Similarly, the gene expression profile of the tumor samples formed two main clusters correlating with the presence of TLSs in histopathological assessment. The tumors with TLSs showed upregulation of genes associated with lymphocyte migration to TLSs. *CCL19*, *CCL21,* and *SELL* are expressed by high endothelial venules (HEVs), critical postcapillary venules evident for both lymphocyte migration to TLSs(26), but also for TLS development(7). Lymphocytes migrate from the periphery to the target organ according to the CCL19 and CCL21 gradient using CCR7(27), after which by upregulation of CXCR5 T and B cells designated for GC reaction, here in TLSs, enter to B cell follicle by following CXCL13 gradient(28,29). As signs of infiltrating B cells, tumors with TLSs also had upregulated CD20 coding *MS4A*(10), and *BTLA* expressed by various B cells and Tfh cells also known to control the GC reaction(30). We also observed upregulation of *FCRL2*(31,32) as a sign of active GC reaction, and *CD27*(33), a memory marker for B cells. Also, *IL7R* seen in activated Th cells(34), *IL2RG* (the common gamma chain)(35), and *CD3D* and *CD3E* evident for T-cell receptor complex(36) where upregulated as a possible sign of general activation of T effector cells related to cancer immunity. Furthermore, we observed that tumors with TLSs contained increased numbers of B, Th, and Tregs, which, together with the gene expression profile, suggest the activation of targeted antibody responses in the TLS, implicating that targeting TLSs could be a promising therapeutic avenue for enhancing antitumor immunity, especially in ovarian cancers expressing CCL19, CCL21, and CXCL13 as biomarkers for TLSs.

Among the distinct clinical groups, we observed enriched GC B cells with enhanced spatial interactions with Tfh cells in the chemo-exposed TLSs as compared to the chemo-naïve TLSs. In general, GC B cells are found from the GCs where they alternate between rounds of somatic hypermutations in the dark zone and competition of surviving signals from Tfh in the light zone. Our results underline how specific the cell neighborhoods are in TLSs, and how significant cell-to-cell connections are for the GCs. Further, the migration of T and B cells to tumor areas leading to the formation of TLSs(26,37) has been associated with CD31+ HEV(38). Consistently, we observed increased frequencies of CD31+ endothelial cells in the chemo-exposed TLSs, potentially indicative of sustained TLS formation and maturation. We observed the enrichment of Tfh, Tfr, and Tfc cells and enhanced spatial interactions between GC B cells and Tfh and Tfr cells in the TLSs of *BRCA1*mut HGSC. These findings together suggest increased follicular T cell accumulation in the TLSs of *BRCA1*mut HGSC regardless of the lack of differences in the total GC B cell counts in the TLSs or the WSIs(4). Further, in line with our findings from WSI overall distinct spatial TMEs in *BRCA*mut HGSC(5), we observed a reduction of CD11c+ APCs, and CD163+ and IBA1+CD163+CD11c+ macrophages particularly in the TLSs of *BRCA1*mut patients. Altogether, the results suggest activated adaptive immunity manifested as TLSs in the chemo-exposed and *BRCA1*mut HGSC, however, the exact mechanisms remain unknown warranting future studies in larger cohorts with patient-matched sample sets.

In addition to the single-cell phenotypes and cellular neighborhoods, emerging evidence suggests that also cellular phenotypes are affected by their spatial context(5,39). In anti-tumor immunity, PD1+ T cells are often considered exhausted T cells, whereas in germinal centers, e.g., of SLOs and TLSs, PD1+ T cells usually are subsets of follicular T cells (Tf). Tfh and Tfr cells are specialized subsets of T helper cells needed in affinity-driven evolution and selection of B cells and are found both in TLSs and GCs characterized mainly by chemokine receptor CXCR5 in addition to canonical T cell markers. Herein, the direct spatial focus on TLSs enabled the phenotypic characterization of the CD3+CD4+PD1+ cells as Tf helper cells (Tfh) and the CD3+CD4+PD1+FOXP3+ as Tf regulator cells (Tfr) even without information on CXCR5 staining. Similarly, both PD1-positive and -negative CD8 T cells were significantly enriched in the T cell neighborhoods and showed avoidance patterns towards cancer and myeloid cells. Of the CD8 cells, PD1-positive cells were adjacency to GC B cells supporting the notion that the CD3+CD8+PD1+ cells in TLSs here represent a follicular subset of CD8 T cells (here stated as Tfc cells) rather than the classical exhausted Tc cells, further strengthening the notion that the phenotypic roles of the cells in the TME are dependent on their spatial context.

In summary, our findings suggest that the presence of TLSs associates with a distinct TME composition and gene expression profile with upregulation of chemokines CCL19, CCL21 and CXCL13. Moreover, we present a detailed spatial composition of TLSs, with enhanced GC functions and adaptive anti-tumor immunity particularly in the chemo-exposed and *BRCA1*mut tumors. We acknowledge that our study is limited in the number of samples especially in the CyCIF cohort, and thus pave the way for future studies in larger patient cohorts. Nevertheless, our findings provide new insights into the spatial biology of TLSs opening new possibilities for precision immunotherapeutic targeting in ovarian cancer.

## List of Abbreviations

HGSC: high-grade serous ovarian cancer
TME: tumor microenvironment
TIL: tumor-infiltrating lymphocyte
TLS: tertiary lymphoid structure
GC: germinal center
Tfh: follicular T helper cell
Tfr: follicular T regulator cell
Tfc: follicular cytotoxic T cell
FFPE: Formalin-fixed paraffin-embedded
CyCIF: cyclic immunofluorescence
APC: antigen-presenting cell
HEV: high endothelial venule

## Declarations

### Ethics approval and consent to participate

This study was conducted in accordance with the Declaration of Helsinki and was approved by the Dana-Farber Cancer Institute institutional review board (DFCI 15-550). All patients provided written informed consent to participate in the study.

### Availability of data and material

All code used in the study are available at https://github.com/SarkkinenJ/OV_CA_TLS. H&E images and CycIF data of TLS crops (TLS images, masks, single-cell data) are available at https://doi.org/10.7303/syn51375062. The CycIF WSI, Nanostring and clinical data are available at https://doi.org/10.7303/syn21569629.

## Authorś contributions

J.S. coordinated the study, and performed cell segmentation, and analyzed all the image data and wrote the manuscript; A.J. performed the Nanostring analysis; J.C. generated the CycIF multiplexed image crops; J.C. and F.P. generated the single-cell quantitated imaging data, J.S., J.C., E.A, and A.S. performed quality control of images; A.L. together with J.S. and A.J. evaluated the histology of H&E tumor sections; J.S., I-M.L, and A.F did the neighborhood analyses, J.S., E.K., and A.F. designed the study, A.F. supervised the study and wrote the manuscript. All authors contributed to the writing and editing of the manuscript.

