## Supplementary Figures for "Single-cell spatial atlas of tertiary lymphoid structures in ovarian cancer"

S1

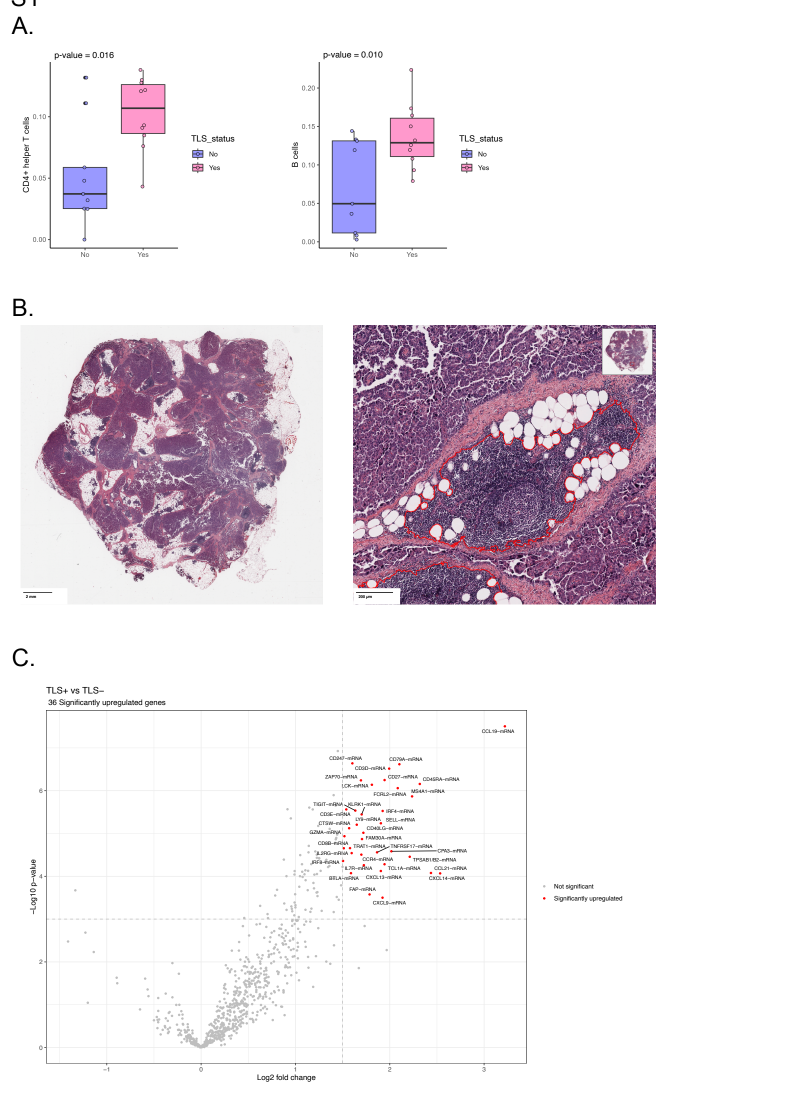

S1

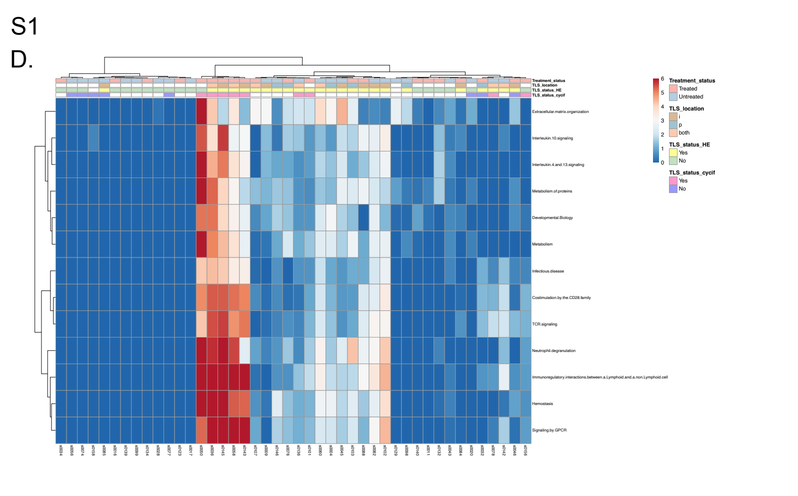

**Supplementary Figure 1: The TLS status reflects cell composition and gene expression profile in TME**
(A) Box and whisker plots showing that ovarian tumors with TLSs have a higher proportion of CD20+ B cells (p = 0.016) and CD4+ T helper cells (CD3+CD4+FOXP3-, p = 0.010) as compared to tumors without TLSs. P-values were calculated with an unpaired Wilcoxon test. (B) An example of H&E stained high-grade serous ovarian cancer sample with TLSs. Left panel shows the whole section is shown, and right panel shows an example of a TLS annotated in red. (C) Volcano plot showing 36 significantly upregulated genes in tumors with TLSs compared to tumors without them. A p-value cutoff of -log10 of 3 and a log2FC of >1.5 or <−1.5 were used. Prior, we filtered for the possible genes, which could have affected by the chemotherapy exposure of the tumor samples (a cutoff of -log10 p-value of 3.0 and a log2FC of >2.5 or <−2.5), however, none of the 36 significantly upregulated genes were affected. (D) Cluster heatmap of KEGG pathway analysis of the differentially expressed genes. A -log10 p-value of 2.5 and a log2FC cutoff of >2.5 or <−2.5 were used to identify significantly upregulated pathways. Clustering was performed with the complete method.

S2

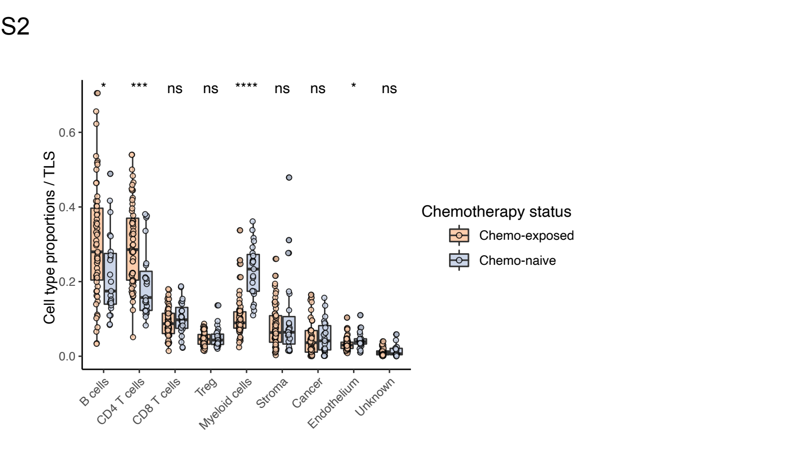


**Supplementary Figure 2:**
**Chemotherapy-exposed TLSs exhibit higher frequencies of myeloid cells**
Box and whisker plots of the frequencies of the main cell lineages within the TLSs. The chemo-exposed TLSs have a smaller proportion of B cells (CD20+; p = 0.01) and CD4 T helper cells (CD3+CD4+FOXP3-; p = 1e-04), and an increased proportion of endothelial cells (CD31+; p = 0.01) and myeloid cells (2e-09). P-values were calculated with an unpaired Wilcoxon test.

S3

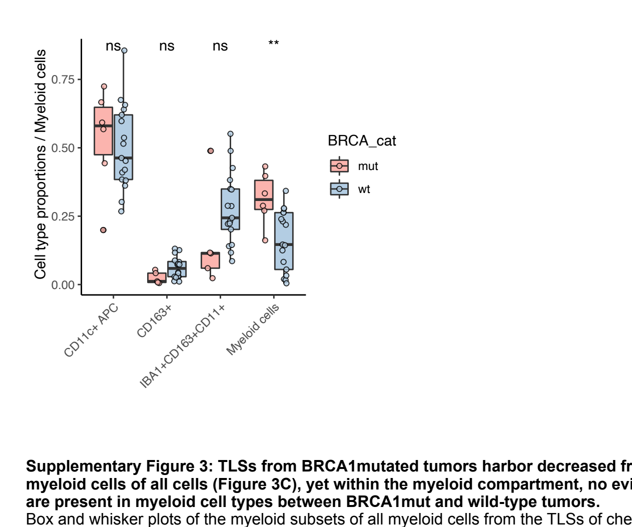


**Supplementary Figure 3: TLSs from BRCA1mutated tumors harbor decreased frequencies of myeloid cells of all cells (Figure 3C), yet within the myeloid compartment, no evident changes are present in myeloid cell types between BRCA1mut and wild-type tumors.**
Box and whisker plots of the myeloid subsets of all myeloid cells from the TLSs of chemo-exposed tumors. Unspecific myeloid cells were decreased (p ≤ 0.01) in the *BRCA1*mut TLSs as compared to the wild-type TLSs, yet no statistical difference was observed in the frequencies of other myeloid subsets. P-values were calculated with an unpaired Wilcoxon test.

S4

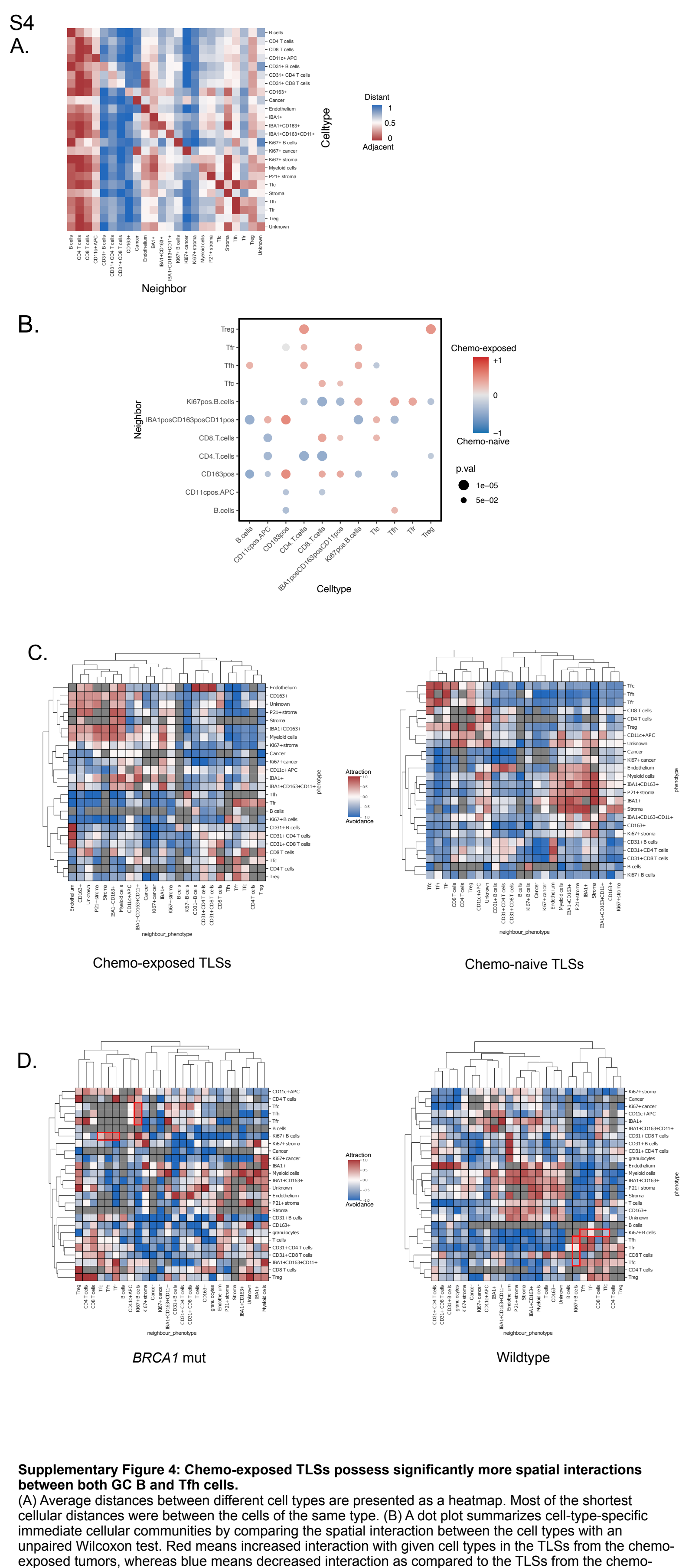


**Supplementary Figure 4: Chemo-exposed TLSs possess significantly more spatial interactions between both GC B and Tfh cells.**
(A) Average distances between different cell types are presented as a heatmap. Most of the shortest cellular distances were between the cells of the same type. (B) A dot plot summarizes cell-type-specific immediate cellular communities by comparing the spatial interaction between the cell types with an unpaired Wilcoxon test. Red means increased interaction with given cell types in the TLSs from the chemo-exposed tumors, whereas blue means decreased interaction as compared to the TLSs from the chemo-naive tumors. The size of the dot reflects statistical significance. (C) Cluster heatmaps show patterns of cell-type-specific immediate cellular communities in TLSs from chemo-exposed and chemo-naive tumors. (D) The cellular communities for *BRCA1*mut and wild-type TLSs are summarised using cluster heatmaps. The red rectangles highligh increased spatial attractions between the GC B and all follicular T cell populations in *BRCA1*mut TLSs as compared to the wild-type TLSs. In C-D, p-values were calculated by subtracting the permuted mean from the observed mean divided by the number of permutations, where the red color reflects an increased attraction with given cell types, whereas the blue color means avoidance compared to random. The 10 nearest neighbors were used for B-D to define immediate cell-type-specific communities for every cell.

S5

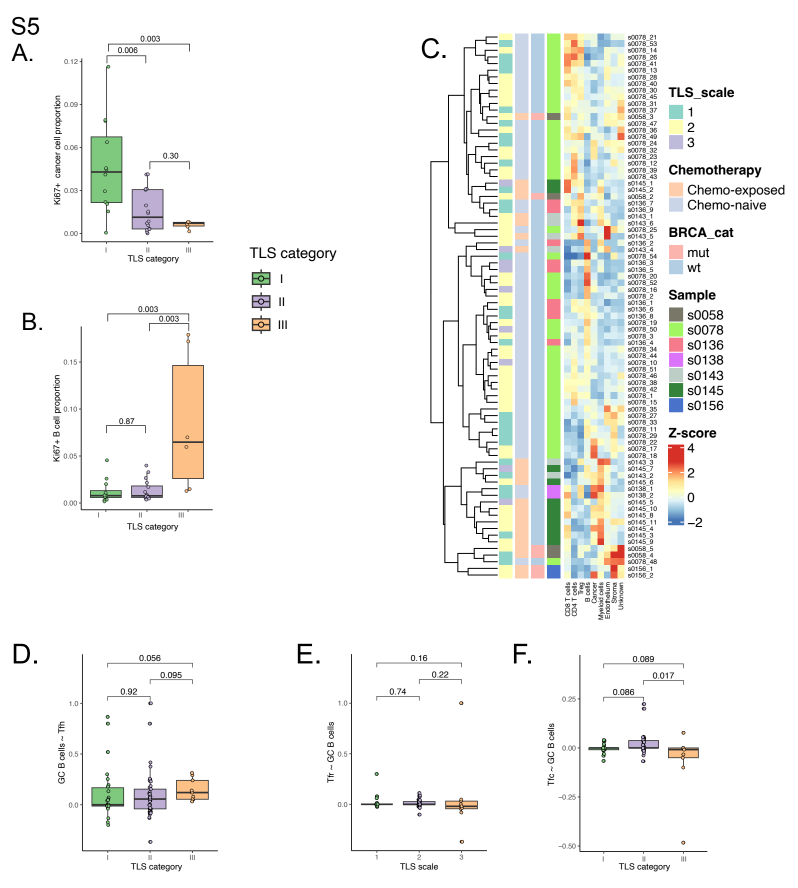


**Supplementary Figure 5: Cell composition and spatial interactions change during the development of TLS**
The proportion of (A) the Ki67+ cancer and (B) GC B cells of all cells in the TLSs with GC (TLS-III) compared to TLS-I and TLS-II. (C) Cluster heatmap of proprotions of main cell lineages in individual TLSs shows heterogeneity between individual TLSs and samples and how TLSs did not clustered along developmental TLS-categories. Clustering was achieved using the complete method. Scaling to z-scores was performed cell lineage-specifically. (D) The GC B cell communities showed a trend of increased interactions with the Tfh cells in TLS-III compared to TLS-I (p = 0.056) and TLS-II (p = 0.095). (E) The GC B cells were neighboring Tfr cell communities equally in all developmental TLS-categories, whereas (F) they were neighboring Tfc cells slightly more in TLSs with B cell follicle lacking a GC than in TLSs with B cell follicle containing a GC (TLS-II vs TLS-III, p = 0.017). In A-B and D-F, p-values were calculated with an unpaired Wilcoxon test.
